# Host circadian rhythms are disrupted during malaria infection in parasite genotype-specific manners

**DOI:** 10.1101/670240

**Authors:** Kimberley F. Prior, Aidan J. O’Donnell, Samuel S. C. Rund, Nicholas J. Savill, Daan R. van der Veen, Sarah E. Reece

**Affiliations:** Institute of Evolutionary Biology & Institute of Immunology and Infection Research, University of Edinburgh, Edinburgh, UK; Department of Biological Sciences, University of Notre Dame, Notre Dame IN, USA; Faculty of Health and Medical Sciences, University of Surrey, Guildford, UK

**Keywords:** Plasmodium, circadian rhythm, circadian clock, sickness behaviour, virulence

## Abstract

Infection can dramatically alter behavioural and physiological traits as hosts become sick and subsequently return to health. Such “sickness behaviours” include disrupted circadian rhythms in both locomotor activity and body temperature. Host sickness behaviours vary in pathogen species-specific manners but the influence of pathogen intraspecific variation is rarely studied. We examine how infection with the murine malaria parasite, *Plasmodium chabaudi*, shapes sickness in terms of parasite genotype-specific effects on host circadian rhythms. We reveal that circadian rhythms in host locomotor activity patterns and body temperature become differentially disrupted and in parasite genotype-specific manners. Locomotor activity and body temperature in combination provide more sensitive measures of health than commonly used virulence metrics for malaria (e.g. anaemia). Moreover, patterns of host disruption cannot be explained simply by variation in replication rate across parasite genotypes or the severity of anaemia each parasite genotype causes. It is well known that disruption to circadian rhythms is associated with non-infectious diseases, including cancer, type 2 diabetes, and obesity. Our results reveal that disruption of host circadian rhythms is a genetically variable virulence trait of pathogens with implications for host health and disease tolerance.

## Background

Circadian rhythms include endogenous and entrainable oscillations in physiology and behaviour with a duration of about 24 hours [1]. Underpinning these rhythms are circadian clocks, which are distributed across the tree of life [2]. Clocks are assumed to be evolutionarily advantageous because they enable their owners to anticipate daily environmental rhythms, enabling organisms to prepare and undertake fitness-determining activities, such as foraging and reproduction, at the most appropriate time of day [3,4]. Additionally, circadian clocks allow for the temporal coordination (or separation) of internal processes [5]. Circadian clocks are reset (entrained) daily by external environmental cues (Zeitgebers), the most prominent including light and time of feeding. In mammals, environmental time-of-day information is received by the suprachiasmatic nuclei (SCN) of the hypothalamus, known as the central clock, via the retinohypothalamic tract. This is transmitted to peripheral clocks in the rest of the body, likely via outputs such as body temperature rhythms and hormone levels. At the same time, non-photic Zeitgebers such as food intake can uncouple peripheral clocks from the SCN and alter their entrained phase [6]. Disruption of the coordination between the SCN and peripheral clocks is a consequence of modern lifestyles, particularly long-term shift work, and associated with a number of non-infectious diseases including cancer, type 2 diabetes, depression and obesity [7,8].

Whilst the association between non-infectious diseases and circadian disruption are well-known, the role that circadian rhythms play in infectious disease has received less attention [9,10]. There are links between infection, inflammation and the SCN [11,12]. Evidence of this affecting circadian rhythms includes: arrhythmic activity patterns in fruit flies infected with bacteria [13], a shortening of the duration (period) of the locomotor activity rhythm in *Trypanosoma*-infected mice [14], and reductions in amplitude of body temperature and locomotor activity rhythms in SIV-infected monkeys [15]. The extent to which circadian rhythm disruption is a cost of being infected, or if it is somehow beneficial to hosts or pathogens is unclear. This is in part due to the complexity caused by interactions between rhythms exhibited by hosts and by pathogens, whom each have different interests during infection. For example, some pathogens may profit from disrupting the circadian rhythms of the host, as suggested for influenza virus whereby interference with clock mechanisms enhances viral replication [16]. Conversely, in cases where the pathogen relies on the host to supply resources from foraging [17,18], or to provide transmission opportunities from interacting with conspecifics [19], pathogens may benefit from bolstering host rhythms. Circadian rhythms in host defences may optimise protection against infection and herbivores, as proposed for Arabidopsis plants, which time their anti-herbivore defences to match the circadian foraging behaviour of caterpillars [20,21]. The extent to which genetic variation amongst pathogens and/or hosts shapes rhythms during infection is unclear. Understanding this would help determine whether altered rhythms are a host response to infections (i.e. a manifestation of pathology or an adaption to control pathogens) and/or are strategically induced by pathogens (i.e. a parasite manipulation). Here, we investigate whether host circadian rhythms are disrupted in a pathogen-genotype dependent manner. We follow infections with three genetically distinct genotypes of the murine malaria parasite, *Plasmodium chabaudi*, and quantify rhythms in a host behavioural trait (locomotor activity) and a physiological trait (body temperature) (see Fig. 1 for experimental design). We expect host rhythms to be altered during infection because lethargy, anorexia and hypothermia are common symptoms of malaria infection [22]. Our parasite genotypes are well characterised and are known to cause different degrees of virulence, in terms of weight loss, anaemia, and mortality risk [23,24,25,26]. We couple the relative ease of tracking parasite dynamics and host rhythms during malaria infection with a new technology to non-invasively monitor locomotor activity and body temperature rhythms throughout complete infection cycles at high temporal resolutions.

**Fig. 1.**
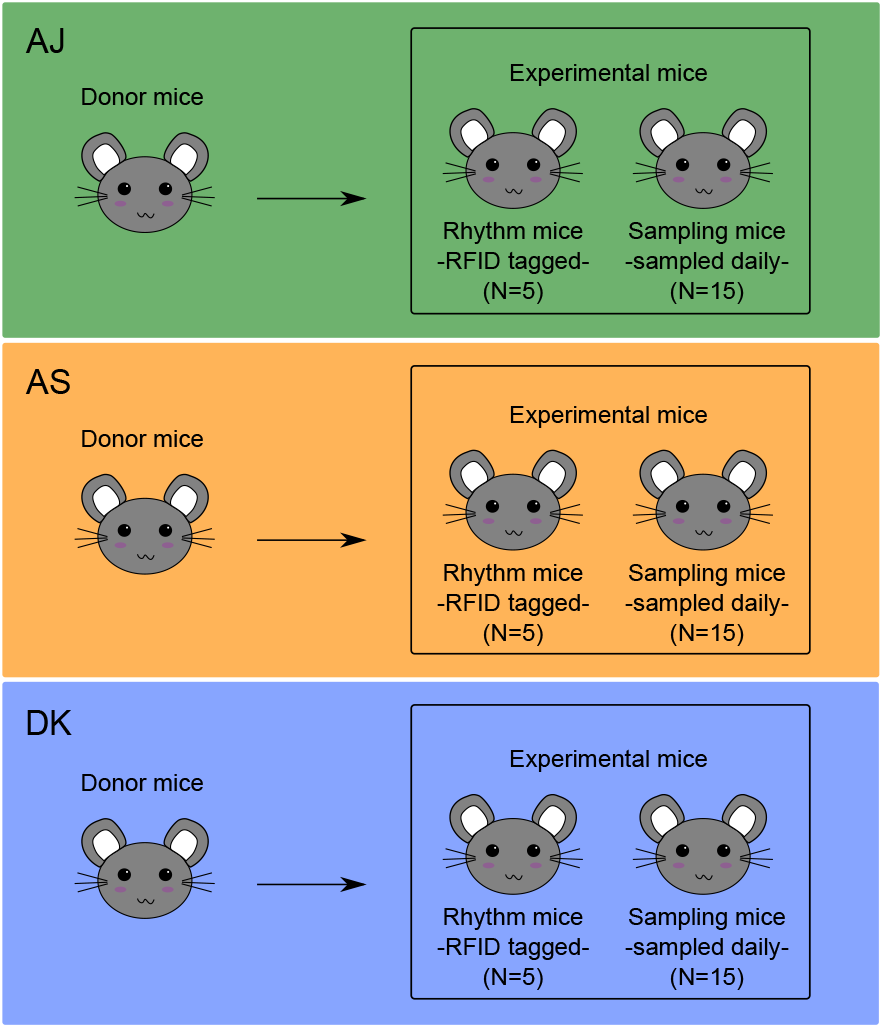
Experimental design. We separately raised three genotypes (AJ, AS, DK) of *Plasmodium chabaudi* in donor mice before inoculating them into 3 groups of 20 experimental mice. We tagged 5 experimental mice per genotype with RFID probes to monitor locomotor activity and body temperature non-invasively (“rhythm mice”) and we blood sampled 15 experimental mice per genotype to monitor parasite and host dynamics once per day (“sampling mice”). We followed host rhythms and infection dynamics throughout 14 days of infection.

Our specific aims are to answer two questions. First, are our measured host circadian rhythms altered in a parasite-genotype dependent manner throughout the entire infection – from the asymptomatic phase, through increasing severity of sickness symptoms, and the return to health? Second, is parasite-genotype dependent disruption to host rhythms associated with their varying levels of virulence and replication rates? We find that: (i) disruption to locomotor activity and body temperature rhythms occurs during infection in a parasite-genotype dependent manner; (ii) virulent genotypes are associated with more severe effects on host rhythms; (iii) locomotor activity and body temperature rhythms can change independently; and (iv) rhythm disruption is only explained in part by variation in the severity of anaemia or parasite density induced by each genotype. We offer hypotheses for why the genotypes differentially affect host rhythms and why locomotor activity and body temperature rhythms might change independently.

## Results

### Host rhythms vary during infection in a parasite genotype-specific manner

Figure 2a illustrates the four segments of infection based on the relationship between anaemia and parasite density: “asymptomatic” (days 3-5, in white), “moderate” (days 6-8, in medium grey), “severe” (days 9-11, in dark grey), and “recovery” (days 12-14, in light grey) (following [27]). Daily rhythms in locomotor activity and body temperature during each segment of infection for hosts infected with each genotype are shown in Fig. 2b.

**Fig. 2.**
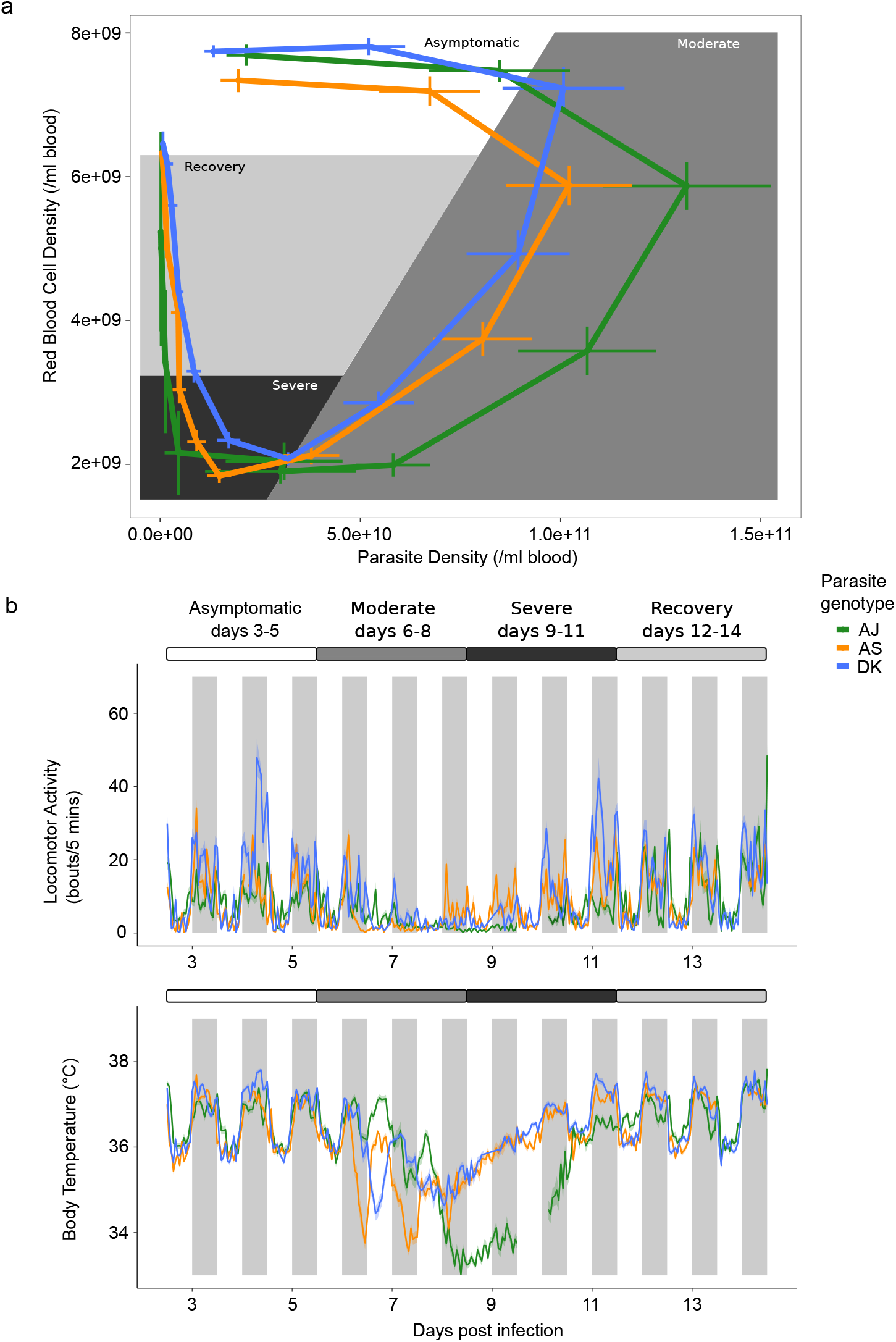
Parasite genotype-specific effects on host sickness, locomotor activity and body temperature rhythms during malaria infection. a) Disease map of host sickness using the relationship between mean ± SEM red blood cell (RBC) and mean ± SEM parasite density (adapted from [27]) for three parasite genotypes (N≤15 per genotype: green=AJ, orange=AS, blue=DK) measured each day post infection (PI) for 14 days. The map falls into 4 three-day segments. (i) Hosts are considered “asymptomatic” (white, days 3-5 PI) until RBC density begins to drop; (ii) Hosts experience “moderate” symptoms (medium grey, days 6-8 PI) until RBC density reaches its minimum; (iii) “severe” symptoms (dark grey, days 9-11 PI) spans the period of extremely low RBC densities; and (iv) hosts are in “recovery” (light grey, days 12-14 PI) until RBC density returns to the level before infection. b) Mean ± SEM hourly locomotor activity and body temperature (see Supplementary Table S1 for further explanation) for 14 days of infection with the same three parasite genotypes (N=5 per genotype: green=AJ, orange=AS, blue=DK). Note in B the line break on day 9-10 PI for the AJ genotype which represents missing data.

#### Across-infection patterns in locomotor activity rhythms

First, we assess whether there is variation between genotypes in host locomotor activity and body temperature rhythms across segments of infections, focussing on metrics for time of peak (specifically, “phase” which is defined as the position of a point in the cycle of a waveform) and amplitude (here measured as change of a variable between peak and trough) [28]. We find differences in locomotor activity patterns induced by parasite genotypes across the different segments of infection, as revealed by significant interactions between sine or cosine terms, parasite genotype and infection segment (Fig. 2, Fig. 3a, sine: χ^2^=17.74_(9,1383)_, p<0.0001, cosine: χ^2^=8.51_(9,1383)_, p<0.0001, R^2^ for model fit = 0.51; see Supplementary Table S2 for effect of genotype). We find that locomotor activity rhythms begin to differ between mice infected with different parasite genotypes during the “asymptomatic” segment: AJ infected mice display reduced amplitude rhythms (amplitude using average model fit: AJ = 5.20 bouts/5 mins; AS = 15.23, DK = 23.07) and a slightly delayed time of peak in locomotor activity in the circadian cycle compared to the other genotypes (the ZT time of peak corresponding to the maximum average model fit: AJ = ZT19.3, AS = ZT17.02, DK = ZT18.4). During the “moderate” segment, AJ infected mice become arrhythmic in locomotor activity whereas AS and DK infected hosts maintain rhythms but lose amplitude (AS = 6.89 bouts/5 mins, DK = 4.51) and the timing of peak locomotor activity advances in the circadian cycle (AS = ZT14.72, DK = ZT17.25). During the “severe” segment, AJ infected mice regain rhythmicity in locomotor activity, although the amplitude remains low (2.01 bouts/5 mins), while the amplitude in locomotor activity increases for AS and DK (AS = 10.64, DK = 18.14). Also, the time of peak locomotor activity for AJ and DK infected mice occurs earlier in the circadian cycle while AS mice returns to the peak time observed in the “asymptomatic” segment (“severe”: AJ = ZT15.41, AS = ZT18.63, DK = ZT15.64). During the “recovery” segment, rhythms of mice infected with AS and DK return to the amplitude (AS = 14.59 bouts/5 mins, DK = 19.12) and time of peak of the “asymptomatic” segment (AS = ZT17.71, DK = ZT17.02). AJ infected mice have rhythms of greater amplitude in locomotor activity during the “recovery” segment (AJ = 15.42) compared to those observed in the “asymptomatic” segment, becoming similar to AS and DK infected mice. Rhythms in locomotor activity of AJ infected mice also have a similar time of peak locomotor activity to AS and DK during the “recovery” segment (AJ = ZT17.48) compared to the “asymptomatic” segment. We then examine how locomotor activity rhythm amplitudes differ in more detail by calculating night-day differences in levels of locomotor activity. Consistent with the previous analysis, we find significant differences in the level of locomotor activity induced by infection with different parasite genotypes across the different segments of infection (interaction between time-of-day, parasite genotype and infection segment: χ^2^=8.99_(6,115)_, p<0.0001, R^2^ for model fit = 0.94; Fig. 3b). To explore this further, we separately analyse night and daytime changes to levels of locomotor activity and observe a significant interaction between genotype and infection segment for both night-time and daytime (night: χ^2^=14.79_(6,58)_, p<0.0001, R^2^ for model fit = 0.91; day: χ^2^=3.50_(6,58)_, p=0.008, R^2^ for model fit = 0.51; see Supplementary Table S3 for effect of genotype). In addition to the changes observed in the phase and amplitude analysis, we also find that: (i) Across all segments and parasite genotypes, mice are on average 3.5-fold more active in the night-time (night-time mean 10.81±0.77) than in the daytime (3.13±0.19), in line with the nocturnality of these mice. (ii) Night-time locomotor activity varies more during infection than daytime locomotor activity (range across all segments, night-time 3-20 locomotor activity bouts/5 mins; daytime 2-5). (iii) Night-time locomotor activity decreases as infections progress. This decrease is largest for AJ infected mice, intermediate for AS infections, and least for DK infections. Conversely, night-time activity is regained sooner for DK and AS infected mice than AJ infected mice (Fig. 3). (iv) Arrhythmicity in AJ infected hosts during the “moderate” and “severe” segments is driven by a loss of night-time activity (Fig. 3).

**Fig 3.**
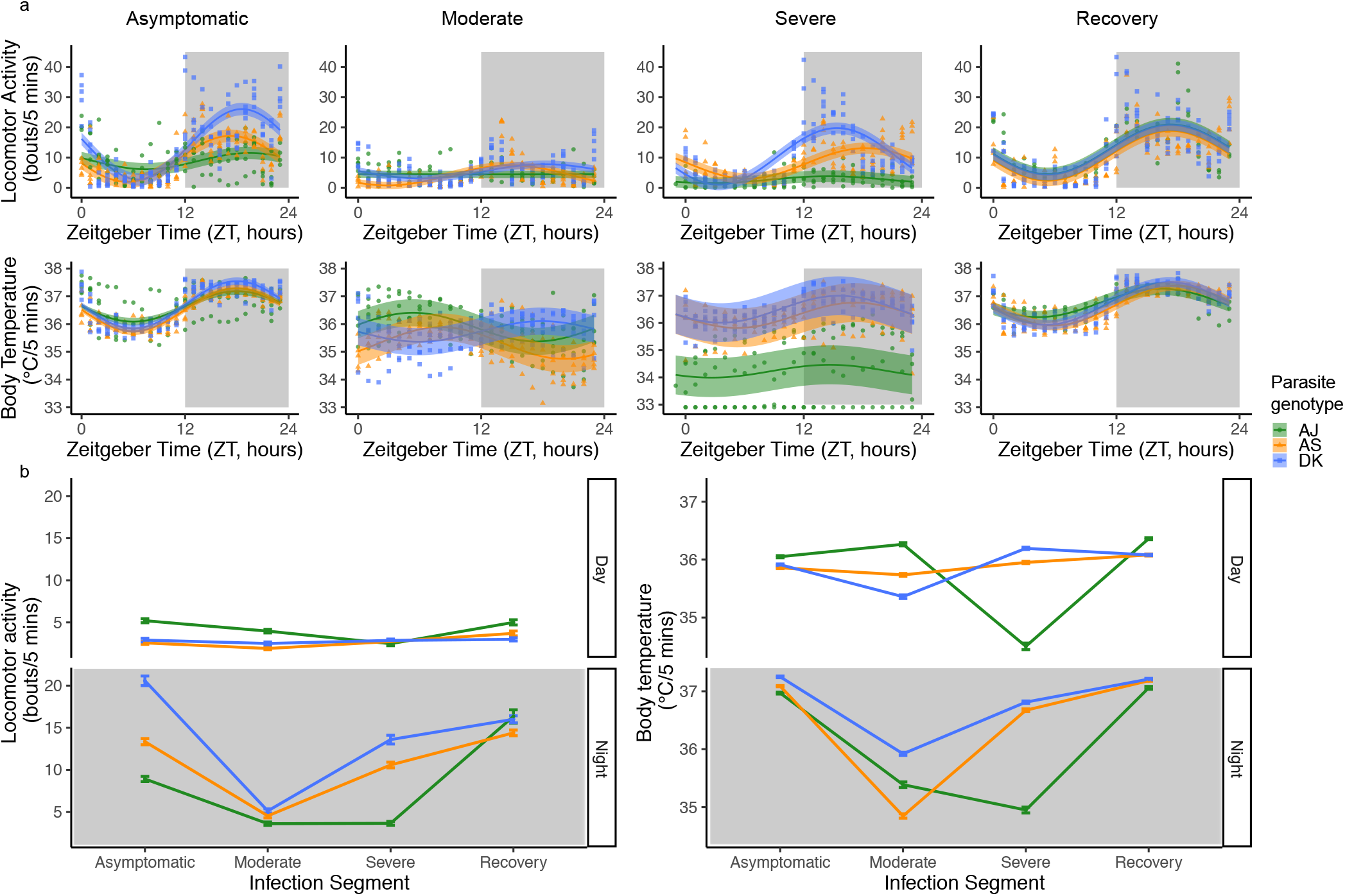
Locomotor activity and body temperature profiles for each 3-day infection segment. a) All data with model predictions (fitted model for the average subject, with 95% confidence intervals calculated by bootstrapping N=500). Each data point behind the model prediction is a 3-day locomotor activity or body temperature average for each mouse (N≤15 per genotype: green circles=AJ, orange triangles=AS, blue squares=DK) at every hour (24-hours in total) and the model is fit to these averaged data points. Time is in Zeitgeber Time (ZT) which is the number of hours since lights on (ZT0) and ZT12 is the start of lights off (as indicated by shaded area). b) Mean ±SEM levels of locomotor activity or body temperature during night time or daytime for the infection segments (N≤5 per genotype: green=AJ, orange=AS, blue=DK). For night-day comparisons we use the average amount of locomotor activity and body temperature for each segment of infection, using ZT14-22 as night time and ZT2-10 as daytime to avoid any effects of dark-light transitions. Night time is indicated by the shaded area. Light and dark bars indicate lights on (7am/ZT0) and lights off (7pm/ZT12).

#### Across-infection patterns in body temperature rhythms

We repeat the above analyses (for locomotor activity) now for body temperature. We find differences in body temperature patterns induced by parasite genotypes across the different segments of infection, as revealed by significant interactions between sine or cosine terms, parasite genotype and infection segment (Fig. 2, Fig. 3a, sine: χ^2^=4.31_(8,1742)_, p<0.0001, cosine: χ^2^=7.06_(8,1742)_, p<0.0001, R^2^ for model fit = 0.53; see Supplementary Table S2 for effect of genotype). During the “asymptomatic” segment, AJ infected mice display lower amplitude rhythms in body temperature compared to infection with AS and DK (amplitude using average model fit: AJ = 1.12°C, AS = 1.54°C, DK = 1.7°C) and all mice have their peak body temperature at a similar time (the ZT corresponding to the maximum average model fit: AJ = ZT18.86, AS = ZT17.94, DK = ZT17.94). During the “moderate” segment, unlike rhythms in locomotor activity, AJ infection does not generate arrhythmicity in body temperature although all genotypes reduce amplitude (AJ = 1.07°C, AS = 1.2°C, DK = 0.72°C). Furthermore, time of peak body temperature advances in the circadian cycle for AJ and AS (AJ = ZT5.98, AS = ZT8.28), but remains similar to the “asymptomatic” segment for DK (DK = ZT17.25). During the “severe” segment, amplitude in body temperature remains dampened (AJ = 0.41°C, AS = 0.95°C, DK = 0.95°C) and the time of peak body temperature becomes closer to that in the “asymptomatic” phase (AJ = ZT15.41, AS = ZT17.71, DK = ZT16.1). In the “recovery” segment, body temperature rhythms remain slightly dampened for all mice (AJ = 1.02°C, AS = 1.42°C, DK = 1.54°C) and peak time returns to match the “asymptomatic” segment (all mice ZT16-18).

Next, we examine how body temperature rhythms differ by calculating night-day differences. We find significant two-way interactions between all explanatory variables (Fig. 3b) – time-of-day: parasite genotype χ^2^=4.15_(2,115)_, p=0.02), time-of-day: infection segment (χ^2^=16.21_(3,115)_, p<0.0001), parasite genotype: infection segment (χ^2^=15.86_(3,115)_, p<0.0001), R^2^ for model fit = 0.75; see Supplementary Table S3 for effect of genotype. As for locomotor activity, we separately analyse night and daytime changes in body temperature and find a significant interaction between genotype and infection segment for both night-time (χ^2^=6.20_(6,58)_, p=0.0002, R^2^ for model fit = 0.78) and daytime (χ^2^=11.12_(6,58)_, p<0.0001, R^2^ for model fit = 0.62). In addition to the patterns revealed by considering amplitude and time of peak, we find: (i) Across all segments and parasite genotypes, mice are on average more than 0.5 of a degree Celsius increase in body temperature in the night-time (night-time mean 36.44°C±0.14) than in the daytime (35.82°C±0.10), following locomotor activity patterns. (ii) Night-time body temperature varies more during infection than daytime body temperature (range across all segments, night-time 34.85°C-37.25°C; daytime 34.50°C-36.36°C). Further, in the daytime, body temperature varies more than locomotor activity, especially for AJ infected mice with genotype differences emerging during the “severe” segment (see significant genotype comparisons in Supplementary Table S3). (iii) Following the loss of night-time activity as infections progress, body temperature also decreases (Fig. 3). In addition, AJ infected mice experience a greater reduction in daytime body temperature than locomotor activity in the “severe” segment, and DK infected mice experience a reduction in daytime body temperature in the “moderate” segment. (iv) Greater differences between infection with the different genotypes are revealed by body temperature compared to locomotor activity in both the “moderate” and “severe” segments (Fig. 3). (v) As for locomotor activity, night-time body temperature levels recover sooner for DK and AS infected mice by increasing and becoming similar compared to AJ infected mice (Fig. 3).

### Parasite virulence and disruption to host rhythms

The previous section reveals that, broadly speaking, host circadian rhythms vary during infections in a parasite genotype-specific manner. Overall, the patterns we observe suggest that: (i) locomotor activity rhythms are more sensitive than body temperature rhythms to parasite genotype in the “asymptomatic” segment. (ii) The most variation between infection with different parasite genotypes is exposed in the “severe” segment, with rhythms in AJ infected hosts diverging from rhythms in AS and DK infected hosts (Fig. 3, Supplementary Tables S2 & S3). (iii) The difference in locomotor activity rhythms between infection with AS and DK disappear after the “asymptomatic” segment, but the effects of AJ on locomotor activity and body temperature rhythms are not eroded until during the “recovery” segment (Supplementary Table S3) and even then, AJ mice still have slightly lower amplitude rhythms, particularly in body temperature. Given that AJ is considered the most virulent of these three genotypes according to measures of anaemia, weight loss, and replication rate [24,25] we investigated whether levels of disruption to rhythms during infection correlates with genotype differences in virulence.

For each host, we regress hourly levels of locomotor activity and body temperature for every day post infection (see Supplementary Table S1 “Change in locomotor activity and body temperature, R^2^”) against the levels of each before infection. This gives us an R^2^ value for each day post infection for every mouse which we use as a metric for rhythm similarity: higher values mean rhythms during infection are more similar to rhythms of healthy animals. We find significant differences in rhythm dynamics between mice infected with different parasite genotypes for locomotor activity (Fig. 4a: genotype by day PI interaction χ^2^=2.93_(2,170)_, p<0.0001) and body temperature (Fig. 4b: genotype by day PI interaction χ^2^=2.19_(22,170)_, p=0.003), see Supplementary Table S4 for effect of genotype. The patterns suggest that across all infected mice, locomotor activity rhythms experience greater overall disruption that locomotor activity rhythms (R^2^ range: locomotor activity 0.08 – 0.41; body temperature 0.16 – 0.72), and locomotor activity rhythms are disrupted for more days during infection than body temperature (locomotor activity 6-11 PI, body temperature 7-8 PI). Genotype comparisons reveal, as suggested by the previous analyses, that AJ associates with the most disruption to locomotor activity and body temperature rhythms, particularly during the beginning of infections (“asymptomatic” segment), and disruption occurs up until the “recovery” segment which is longer than the other genotypes (Fig. 3 & 4). AS and DK cause very similar levels of disruption; changes to locomotor activity rhythms occurs earlier in infections with AS than DK but recovery occurs at similar rates.

**Fig. 4.**
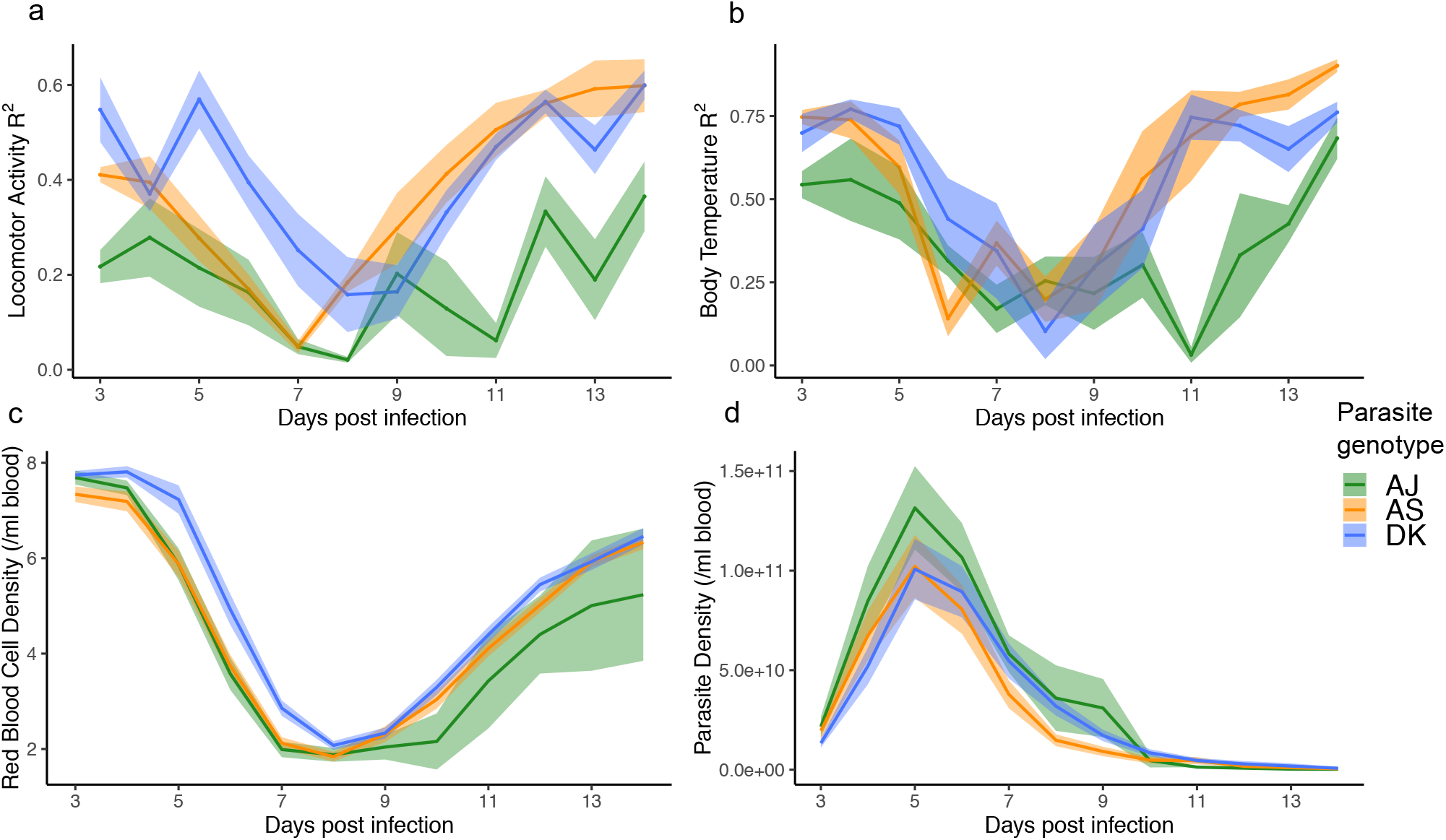
Daily dynamics for disruption to locomotor activity and body temperature rhythms and metrics for parasite virulence. a) Similarity between locomotor activity rhythms on each day post infection compared to before infection (higher R^2^ = rhythms more similar to before infection). b) Similarity between body temperature rhythms on each day post infection compared to before infection. c) Host red blood cell density (anaemia) during infections. d) Asexual parasite density during infections. Mean ± SEM plotted (N≤15 per genotype: green=AJ, orange=AS, blue=DK). Mice were sampled once per day between days 3-14 post infection in c and d. N≤5 per genotype for a and b, and N≤15 for c and d.

#### Genotype differences in parasite density and anaemia do not correlate with disruption to host rhythms

Next, we verify that our genotypes vary in virulence as expected (AJ is the most virulent and DK the least) according to the traditionally used virulence measures of host anaemia and parasite density [24,25]. We find significant differences between mice infected with different genotypes in the densities of red blood cells (Fig. 4c: genotype by day PI interaction χ^2^=2.28_(22,451)_, p=0.001, R^2^ of model fit = 0.90) and parasites (Fig. 4d: main effect genotype χ^2^=14.32_(2,441)_, p<0.0001, main effect day PI χ^2^=51.90 _(11,441)_, p<0.0001, R^2^ of model fit = 0.62) during infections, see Supplementary Table S4 for genotype comparisons. The expected virulence rank is supported by AS and DK infected mice losing RBC at a slower rate than AJ, and AJ infected mice showing the slowest recovery from anaemia. Similarly, AJ infected mice harbour the most parasites throughout infections.

Given the genotype differences in host anaemia and parasite replication, and that most disruption to locomotor activity and body temperature rhythms coincides with minimum RBC density and follows peak parasite density (Fig. 4; days 6-9 PI, see Supplementary Figure 1 for these data plotted as a disease map), we investigated how well RBC loss or parasite density *per se* correlates with rhythm disruption compared to other unknown genotype-specific factors. To do this, we regress the daily measures of rhythm disruptions for each host, as captured by our rhythm similarity metric (see Supplementary Table S1 “Change in locomotor activity and body temperature, R^2^”) for locomotor activity and body temperature against the mean daily measures for RBC and parasite densities. See Supplementary Table S5 for effects of genotype. For locomotor activity disruption, we find an interaction between genotype and RBC density (Fig. 5a, χ^2^=3.55_(2,170)_, p=0.03, R^2^ of model fit = 0.61) as well as an interaction between genotype and parasite density (Fig. 5b, χ^2^=12.00_(2,170)_, p<0.0001, R^2^ of model fit = 0.46). These interactions are driven by AJ causing more disruption to locomotor activity rhythms at the same RBC and parasite density, especially at low RBC and high parasite densities, and by AS causing more severe disruptions at high parasite densities. For body temperature disruption, there is no significant interaction between genotype and RBC density (Fig. 5c, χ^2^=2.56_(2,170)_, p=0.08) but there are significant main effects of genotype (χ^2^=7.66_(2,170)_, p=0.0007) and RBC density (χ^2^=149.19_(1,170)_, p<0.0001), R^2^ of model fit = 0.53. Body temperature rhythm disruption increases as hosts become more anaemic and for a given RBC density, disruption is greater during AJ than AS infection, with DK causing intermediate levels. As for locomotor activity rhythms, we also find that AJ causes greater body temperature rhythm disruption at lower parasite densities than AS and DK which cause similar amounts of disruption (interaction between genotype and parasite density, χ^2^=9.66_(2,170)_, p=0.0001; Fig. 5d, R^2^ of model fit = 0.33). That genotype remains in all of these models (as a main effect or in an interaction) suggests that in addition to the levels of anaemia they cause and their replication rates, other factors inherent to the genotypes must influence host rhythms.

**Fig. 5.**
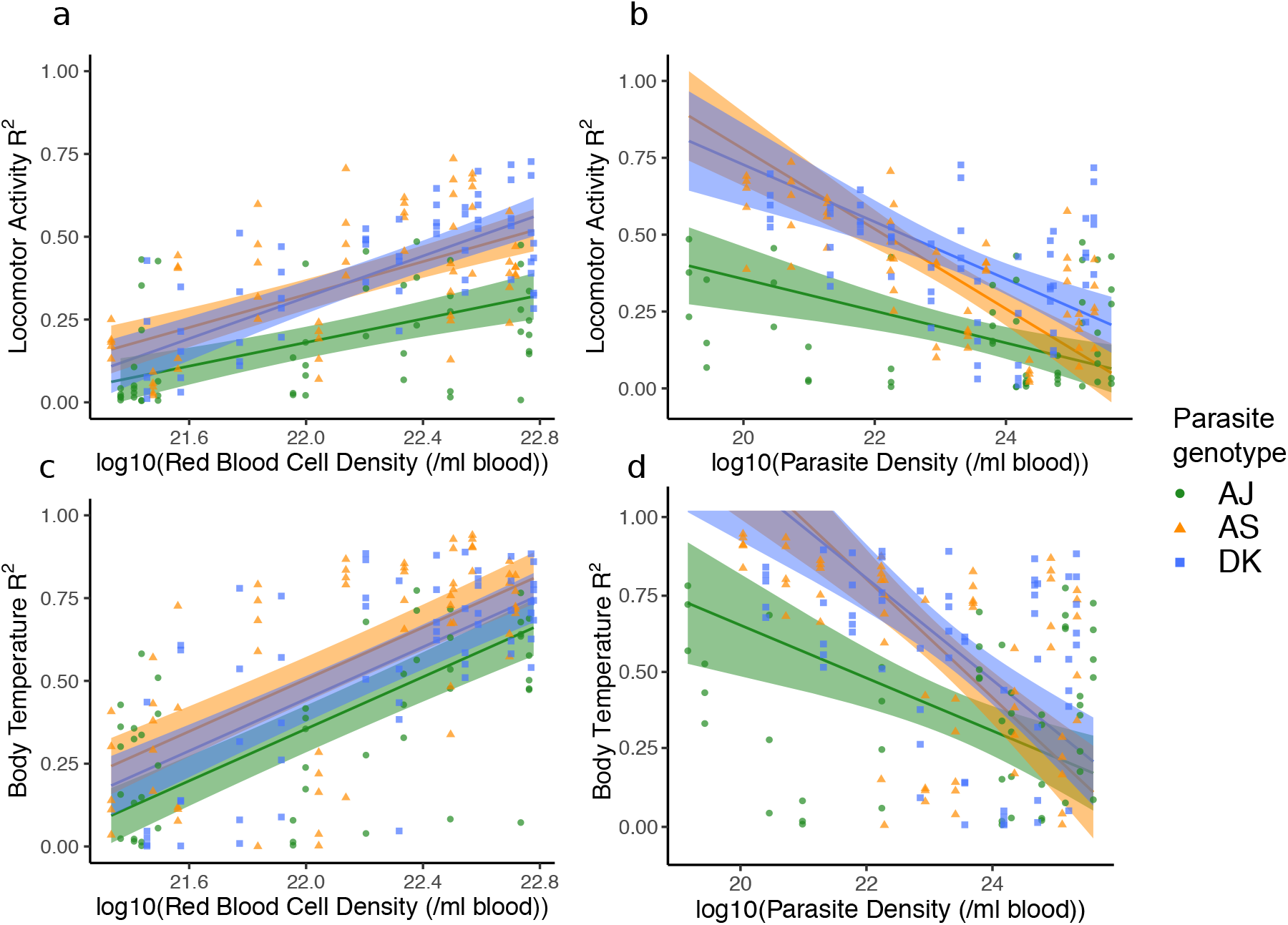
Correlations between red blood cell (RBC) density and parasite density with levels of disruption to locomotor activity (a, b) and body temperature (c, d) rhythms. Hosts infected with each of our parasite genotypes are plotted (green circles=AJ, orange triangles=AS, blue squares=DK), with one measure per day for either red blood cell density or parasite density on the x-axis (calculated from mean of 3 “sampling mice” each day post infection), against all R^2^ values on the y-axis, on each corresponding day post infection (calculated from “rhythm mice”, ≤5 per parasite genotype). A low R^2^ value indicates the pattern of host rhythms is different to the rhythm observed before hosts were infected. RBC and parasite density data are transformed by log10 in the models to improve residual homogeneity. Model predictions are plotted as a solid line for the overall model fit (fitted model for the average subject), with 95% confidence intervals calculated by bootstrapping N=500. Models and error are bounded at 1 on the y-axis, as R^2^ does not go above 1.

## Discussion

We reveal that host rhythms in locomotor activity and body temperature become disrupted during *Plasmodium chabaudi* infections, in a parasite genotype specific manner. The most virulent genotype (AJ) disrupted host rhythms to the greatest extent and for the longest time period during infections, and differences between genotypes become more apparent as hosts progress through infections. However, we unexpectedly find that genotype-specific disruption to rhythms is not associated with genotype differences in parasite density or the degree of host anaemia, revealing a role for intra-specific pathogen variation in shaping the host response to infection. We also find that locomotor activity patterns and body temperature rhythms show changes independently of one another. As expected, we find that infecting mice with *Plasmodium* parasites reduces locomotor activity (lethargy) and body temperature (hypothermia) [22]. We find that night-time locomotor activity and body temperature are indeed being reduced due to infection, but that daytime (when mice are resting) locomotor activity and body temperature are generally not changed as much (Fig. 2 & 3). For locomotor activity patterns, this is likely due to inactivity, which cannot decrease into negative activity, and for body temperature - apart for AJ during the “severe” segment -this may be due to minimum temperature requirements for survival [29,30]. While infection with all parasite genotypes follow broadly similar dynamics, a more detailed dissection of daily rhythms reveals differences between genotypes. For example, the largest locomotor activity differences between genotypes occur during the “asymptomatic” and “severe” segments. DK infected mice are more active during the “asymptomatic” segment compared to AJ and AS infected mice, with >2-fold higher night time locomotor activity compared to AJ infected mice, with AS infected mice intermediate. During the “severe” segment DK infected mice have 4-fold higher night-time locomotor activity compared to AJ infected mice, which remain at low locomotor activity levels as in the “moderate” segment for all genotypes, while AS infected mice increase locomotor activity by a similar amount as DK infected mice. For locomotor activity there are few differences between genotypes during the daytime. For body temperature, most genotype differences occur during the “moderate” and “severe” segments, again with DK infected mice having greater night-time body temperature during these segments compared to AJ infected mice, while body temperature is similar in DK and AS infected mice during the “severe” segment. Daytime body temperature varies between genotypes during the “severe” segment in which AJ infected mice have lower body temperature compared to AS and DK infected mice.

Generally, we find the biggest differences in host circadian rhythms after infection with different parasite genotypes emerging in the “moderate” and “severe” segments of infection, which intriguingly coincides with the development of adaptive immunity [31]. Adaptive immunity is temperature sensitive, with, for example, heat enhancing the rate of lymphocyte trafficking as well as increasing CD8 T cell differentiation towards cytotoxic function and interferon-gamma production [32]. Thus, differences in the degree of hypothermia caused by the parasite genotypes may create variation in the efficiency of the adaptive immune response and subsequently affect recovery rate, potentially contributing to the differences between genotypes illustrated in Fig. 5. However, experiments that independently perturb parasite density and RBC density (for example by using phenylhydrazine to create anaemia) independently of infection are needed to resolve their influences on rhythm disruption by each parasite genotype. Alternatively, in addition to direct parasite-mediated exploitation of host resources, parasite virulence also includes damage due to infection-induced immunopathology, caused by excessive levels of proinflammatory cytokines [33,34]. As the most virulent genotype, AJ may induce more inflammation than the other genotypes and so, hosts take longer to recover from the collateral damage. To test these ideas, animals could be kept on heat pads to maintain body temperature rhythms and explore the role of temperature sensitivity in the immune response, locomotor activity rhythms and other symptoms. Additionally, the role of different or overactive immune responses on circadian rhythms could be tested by blocking or enhancing innate immune effectors, or giving animals live attenuated vaccines [35] or low doses of corticosteroids to dampen immunity.

We find that disruption to locomotor activity and body temperature rhythms do not mirror each other. For example, across infection with all genotypes, locomotor activity rhythms become disrupted sooner and for longer (Fig. 4a and 4b, Supplementary Table S4), and AJ infected mice lose rhythms in locomotor activity during the “moderate” segment but retain rhythmicity in body temperature (Fig. 3). In addition to disruption of locomotor activity and body temperature rhythms occurring at different points in the infection, the nature of changes to their rhythms differs. For example, there is a body temperature rhythm inversion during infections with AJ and AS genotypes which does not happen to locomotor activity rhythms (Fig. 3a, Supplementary Table S2). Changes occur in the timing of peak locomotor activity during sickness but not to the same degree as for body temperature rhythms and there is a reduction in locomotor activity and body temperature rhythm amplitude which occurs earlier in locomotor activity. Behavioural and body temperature rhythms are usually in phase with each other (have similar peak times), with the energetic requirements of physical activity meaning that usually body temperature is also increased. These rhythms are coordinated with the external environment, and while they share a common pacemaker in the SCN they may be differentially regulated [36]. Thus, independent disruption of locomotor activity and body temperature rhythms is unusual and tends to only be seen under an experimentally-induced “forced desynchrony protocols” whereby sleep-wake and body temperature rhythms oscillate with different periods [37,38]. Core body temperature oscillates in a circadian manner and is affected by the basal metabolic rate which is maintained via organism energy expenditure [39]. If sickness suppresses appetite, and therefore reduces the amount of energy intake, this may cause disruptions to metabolism which has a knock-on effect to body temperature. Thus despite being active, sickness may erode the correlation between locomotor activity and body temperature. Furthermore, perhaps body temperature rhythms are inherently more robust due to their role in synchronising peripheral oscillators in organs and tissues (amongst other methods including autonomic innervation, glucocorticoids and feeding rhythms) [40] and so, can be maintained during illness to a greater extent than locomotor activity rhythms. However, if this is the case then it is surprising that once affected, body temperature rhythms experience greater disruption than locomotor activity rhythms.

Whether disruption to rhythms reflects disruption to the circadian oscillator is unclear. This could happen via compromising the operation of the central clock in the SCN, or via interfering with peripheral clocks which then feedback to the SCN clock [40]. Perhaps the sensitivity or perception of Zeitgebers is affected or perhaps mice are constrained by lack of resources (due to anorexia as well as parasite infection) and are unable to follow the instructions provided by their clocks. During infection, host innate immune cells release cytokines (primarily IL-1, IL-6 and TNFα) which act on the hypothalamus in the brain and induces a fever response [32]. The primary circadian oscillator is also located in the suprachiasmatic nucleus within the hypothalamus and whilst its time-keeping should be protected from fever (due to temperature compensation of circadian clocks [41]), it might be sensitive to cytokines [11,12,42]. In support of this notion, simulating an infection can be enough to change the timing of clocks: a lengthening of circadian period and reduced amplitude occurs in cellular rhythms in distal uninfected tissues when Arabidopsis is single-leaf infected with the bacteria *Pseudomonas syringae* or when treated with the plant defence phytohormone salicyclic acid [43]. Furthermore, trypanosome infection in mice shortens the period of circadian clock gene expression in the liver, hypothalamus and adipose tissue [14].

Infection causes pathologies which have negative consequences for host fitness, but there is increasing recognition that some signs of infection are better defined as “sickness behaviours” that promote host survival rather than manifestations of pathology [44]. Generally, sickness behaviours could help individuals cope with the burden of infection by concentrating limited resources on immune functions or by somehow restricting pathogen growth/reproduction. Both hyper- and hypo-thermia are common responses to infection and promote survival [32]. For example, despite a 1°C increase in core body temperature requiring a 10-12% increase in metabolic rate [45], inhibiting fever in these cases increases mortality [46]. Whereas in instances of severe inflammation, lowering temperature promotes survival [47,48]. With acute malaria infection inducing inflammation [49], hypothermia could be best achieved by reducing the amplitude of body temperature rhythms, and energy expenditure could be limited by reducing locomotor activity amplitude. Furthermore, hosts may need to disrupt their core rhythms to override the circadian rhythms of innate immune cells [50], ensuring they can mount an acute response at any time of day. In addition, by disrupting their own rhythms, hosts might limit resources or time-cues required by parasites [17], making parasites more susceptible to clearance by the host [51]. That there is genetic variation amongst parasites in circadian rhythm disruption of the host that correlates with other genetically encoded virulence traits, suggests it is a cost of virulence. If host rhythm disruption is bad for parasites then parasites are likely to be under selection to reinforce host rhythms in physiology and behaviour. Alternatively, disrupting host rhythms may be beneficial to parasites [24,25]: more virulent genotypes tend to be better competitors and inducing greater levels of host disruption may disadvantage other coinfecting genotypes.

In summary, we reveal that malaria infection disrupts host circadian rhythms in behaviour and physiology in parasite genotype dependent manners. It has been proposed that, unlike trypanosome infection, malaria parasites do not affect the circadian rhythms of their hosts [14]. However, this study summarised rhythms over the infection as a whole, but our data reveal short-term rhythm disruption that could easily be missed when summarising an entire infection. While Rijo-Ferreira et al [14] suggest clock gene expression is disrupted in adipose tissue during malaria infection, we have not investigated whether malaria parasites disrupt host rhythms by affecting the working of host circadian clocks, but our data suggest that such studies should focus on the symptomatic phases of infections with virulent genotypes. We propose that circadian rhythm disruption is a more appropriate metric of health than anaemia as it gives an overall systemic view of host physiology and behaviour during sickness, especially during the early phases of infection, and is sensitive to parasite genotype. Whilst we only consider three parasite genotypes, our results suggest host rhythm disruption is a genetically variable parasite trait and cannot be explained by traditional measures of virulence (anaemia and parasite replication rates, Fig. 5). If so, then it may be under selection but it is not clear whether parasites benefit from disrupting or protecting host rhythms. Whether perturbations of immune responses, drug treatment, or genetic diversity of infections increases the exposure of parasite genetic variation to selection is tractable and relevant given the reduction in malaria prevalence and the association between parasite rhythms and tolerance to antimalarial drugs. Our results highlight the potential for circadian rhythms as an arena for host-parasite coevolution. Analysing the costs and benefits of disrupted host rhythms for both hosts and parasites will help reveal to what extent manipulation of their own and each other’s rhythms is involved in parasite offence and host defence.

## Methods

### Infections and experimental design

We used 8-week-old male MF1 mice (bred in-house, University of Edinburgh) with *ad libitum* access to food and drinking water (supplemented with 0.05% para-amino benzoic acid to facilitate parasite growth). We entrained sixty experimental mice to a 12:12h (lights on 7am/ZT0, lights off 7pm/ZT12; ZT is Zeitgeber Time which is the number of hours after lights on) light:dark cycle for 2 weeks prior to, and during, the experiment. Mice were randomly allocated to 3 treatment groups of 20 mice each (see Fig. 1 for experimental design). We intravenously infected all mice with 10^7^ *Plasmodium chabaudi* parasitized red blood cells (RBC) at ring stage. Parasitised RBCs were harvested from donor mice that were on the same light:dark cycle as the experimental mice. Mice in each treatment group received either genotype AJ, AS, or DK (AJ is a more virulent parasite with infections generating greater amounts of anaemia and reaching higher parasite densities, while DK is a less virulent parasite and AS intermediate). We designated 5 mice in each group to the monitoring of rhythms (“rhythm mice”) and they received subcutaneous RFID tags [BioTherm13 RFID (radio-frequency identification) (Biomark, Idaho, USA)] 7 days before infection to continuously record locomotor activity and body temperature rhythms in conjunction with a Home Cage Analysis system (Actual HCA, Actual Analytics Ltd, Edinburgh, Scotland; see Supplementary Methods). The “rhythm mice” remained unsampled throughout the experiment. We designated the remaining 15 mice per group to the monitoring of infection dynamics (“sampling mice”). The “sampling mice” were blood sampled daily (see Supplementary Methods) to quantify parasite dynamics and blood parameters. By separately housing the RFID tagged “rhythm mice” and the blood sampled “sampling mice”, we minimised the effects of disturbance on data collected to examine locomotor activity and body temperature rhythms of the RFID tagged mice. The 15 “sampling mice” were infected in two separate blocks of 10 mice and 5 mice with identical set up to augment sample size in the event of mortality induced by the more virulent genotypes. Data from “rhythm mice” and “sampling mice” were collected between days 3 and 14 post infection (PI). See Supplementary Methods for more information on sampling and data collection.

### Data analyses

We used R v. 3.5.0 (R Core Team 2013) to fit linear models (lmer in the R package “lme4”) and derived minimum adequate models through stepwise model simplification based on Pearson’s chi-square test (using the drop1 function in the R package “stats”). The residual distribution of the model fits were checked visually for normality. We report R^2^ values for each model as a measure of how close the data fit to the regression line (i.e. the variance accounted for by the model). We used mixed-effects models due to repeated measures taken from the same mice. The first section describes how we test for genotype-specific disruption to host rhythms. The second section explains how we verify the genotypes vary in virulence as expected. The final section explains how we determine if variation in disruption to host rhythms is associated with known differences in virulence and replication rate across the parasite genotypes.

#### Genotype-specific effects on host rhythms during infection

Following Torres et al [27] we first split the infection into four equal length segments (“asymptomatic”, “moderate”, “severe”, “recovery”) based on red blood cell and parasite dynamics (see Fig. 2 and Supplementary Table S1 “segment summary”). These segments also reflect the general signs of infection observed during sampling (e.g. lethargy, piloerection). This generates a “disease map” which we use to generalise patterns of host rhythm disruption during different areas of infection in terms of temporal variation in disease parameters. We then ask whether patterns for host locomotor activity and body temperature differ between parasite genotypes, both within and between infection segments. We fit models using either host locomotor activity or body temperature as the response variable. For explanatory variables, we fit parasite genotype, segment of infection, and sine and cosine terms as fixed effects with interactions, with mouse identity as a random effect. We say locomotor activity or body temperature has become arrhythmic (i.e. no detectable rhythm) when both the sine and cosine terms can be removed from the model. We calculate time of peak and amplitude for each rhythm in each rhythmic segment and compare them between parasite genotypes (see Supplementary Table S1 “time of peak” and “amplitude”, [28]). Exploring the differences between genotypes is not computationally possible using this modelling approach because the wave form parameters (sine and cosine terms) are not compatible with good practice for multiple testing of general linear models. Therefore, we only report the R^2^ and AIC values for the effect of genotype (Supplementary Table S2).

Next, we investigate diel variation in host rhythms in more detail by calculating summary variables of locomotor activity and body temperature for night and day (see Supplementary Table S1 “amount of locomotor activity and mean body temperature”). We calculate the mean locomotor activity and body temperature values during each 3-day segment of infection for 8 hours during the night (ZT14-ZT22) and day (ZT2-ZT10) for each mouse. The central 8 hours are used to avoid fluctuations during the light transitions. We again compare the effect of different parasite genotypes on locomotor activity and body temperature within and between the four infection segments. Our models use the summary variables for either host locomotor activity or body temperature as the response variable and, as explanatory variables, parasite genotype, segment of infection and time-of-day (night, day) with interactions, and host identity as a random effect. We use the glht function in the “multcomp” R package to perform pair-wise comparisons between genotypes with Tukey HSD (correcting p values for multiple comparisons) (Supplementary Table S3).

#### Parasite virulence shapes host rhythm disruption

To examine whether the parasite genotypes vary in the patterns of host rhythm disruption during infections, we model either the change in host locomotor activity or body temperature (see Supplementary Table S1; “change in locomotor activity and body temperature, R^2^”) as response variables and day post infection and parasite genotype as explanatory variables, including interactions, and mouse identity as a random effect. Then, to examine whether the parasite genotypes vary in replication rate and in the anaemia they cause, we model either RBC or parasite density as response variables with day post infection and parasite genotype as explanatory variables, including interactions, and mouse identity as a random effect.

#### Replication rate and anaemia explaining disruption to host rhythms

Given that the genotypes differ across infections in host locomotor activity and body temperature rhythm disruption and reach different parasite densities and cause different levels of anaemia, we ask whether rhythm disruption is associated with the densities of RBCs or parasites. We fit models using change in host locomotor activity or body temperature (see Supplementary Table S1 “change in locomotor activity and body temperature, R^2^”) as response variables and parasite genotype, log base10 RBC or parasite density as explanatory variables with interactions, and mouse identity as a random effect. Note, because our design used different mice to monitor rhythms and infection dynamics, the models included the average RBC or parasite density per day as determined by the “sampling mice” (N≤15 per genotype), and locomotor activity and body temperature measures per day from each of the “rhythm mice” (N≤5 per genotype). Again, post-hoc comparisons between genotypes were made with Tukey HSD (Supplementary Table S5).

### Ethics

All procedures were carried out in accordance with the UK Home Office regulations (Animals Scientific Procedures Act 1986; project licence number 70/8546) and approved by the University of Edinburgh.

## Acknowledgements

We thank Petra Schneider, Luke McNally, Pedro Vale and Hannes Becher for advice and discussion. We also thank David Schneider and another anonymous reviewer for their insightful comments.

## Authors’ contributions

KFP, AJOD and SER designed the study; KFP and DvdV performed the data analysis; NJS, SSCR and SER provided project supervision and aided in interpretation of the data; KFP wrote the first manuscript draft. All authors gave final approval for publication.

