## Supplementary Information for "Host circadian rhythms are disrupted during malaria infection in parasite genotype-specific manners"

### Supplementary Methods

The Home Cage Analysis System uses an RFID reader baseplate with antennae located at even spaced positions on the plate, with an electromagnetic field around each antenna. This enables the position and body temperature of each RFID tagged mouse, "rhythm mouse", in the cage to be recorded 0.05 seconds non-invasively, in conjunction with mouse identity. The surface area is 612 × 435 mm with 12 evenly spaced antennae. The position data are converted into an activity transition, here termed "bout", when mice move between antennae.
Mice used to examine parasite dynamics and blood parameters ("sampling mice") were sampled daily at 9am/ZT2 by removing 4μl blood from the tail vein to: (1) measure red blood cell density (RBC; flow cytometry, Beckman Coulter), (2) estimate blood glucose concentration (Accu-Chek, Roche Diagnostics), and (3) make thin blood smears to calculate the proportion of infected RBC (parasitaemia). Rhythms were recorded, and mice blood sampled, for 14 days, until parasites were no longer detectable by microscopy.

### Supplementary Figures

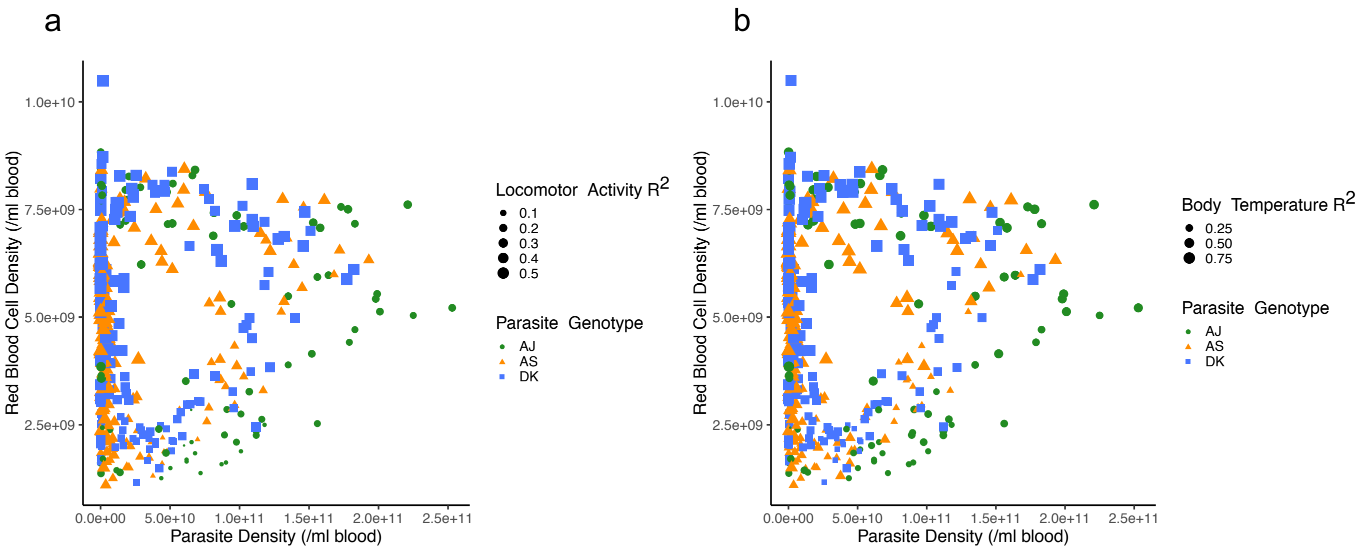

Supplementary Figure S1. Incorporating levels of disruption to locomotor activity (a) and body temperature (b) with parasite genotype-specific effects on host sickness during malaria infection. Disease map of host sickness using the relationship between red blood cell and parasite density for three parasite genotypes (N≤15 per genotype: green circles=AJ, orange triangles=AS, blue squares=DK) measured each day post infection (PI) for 14 days. Size of points corresponds to the similarity between a) locomotor activity rhythms and b) body temperature rhythms on each day post infection compared to before infection (higher $R^2$/larger points illustrates rhythms that more similar to those before infection). Associated with Figures 2, 4 and 5.

### Supplementary Tables

| | Explanation |
|---|---|
| **Segment summary**; Methods:Data analyses:Genotype-specific effects on host rhythms during infection | Infection segments include “asymptomatic”, “moderate”, “severe” and “recovery”. We collapsed data collected during each 3-day infection segment into one 24-hour period by calculating the average locomotor activity and body temperature for every hour during the circadian cycle, for each mouse (i.e. 24 data points for each mouse per segment comprising of the hourly average over 72 hours of infection). |
| **Time of peak**; Methods:Data analyses:Genotype-specific effects on host rhythms during infection | Using the hourly binned data, we fit sine and cosine terms in a linear mixed effects model to the segment summary data. By finding the maximum of the fitted curve we get an estimate for what time of day (ZT) the maximum amount of locomotor activity or body temperature occurs during each infection segment (i.e. the position of a point in time on a wave form). If neither sine nor cosine term are significant in the model, then the data is not deemed rhythmic and neither time of peak nor amplitude are calculated. |
| **Amplitude**; Methods:Data analyses:Genotype-specific effects on host rhythms during infection | By fitting sine and cosine terms in a linear mixed effects model to the segment summary data, the fitted curve provides estimates for the range or peak-to-peak amplitude of the data (distance between maximum and minimum) during each infection segment (i.e. measure of change of a periodic variable over a single time period). |
| **Amount of locomotor activity and mean body temperature**; Methods:Data analyses:Genotype-specific effects on host rhythms during infection | Using the number of activity “bouts” and mean body temperature in 5-minute intervals, we take measures of day and night time locomotor activity using the middle 8 hours of night and day (ZT14-22 and ZT2-10), to remove fluctuations caused by lights off/on. This gives us a measure of the actual amount of locomotor activity and body temperature to |

| | |
|---|---|
| | compare between genotypes (alongside peak phase and amplitude). |
| **Change in locomotor activity and body temperature, $R^2$**; Methods:Data analyses:Replication rate and anaemia explaining disruption to host rhythms | Changes in host rhythms were measured by using daily $R^2$ values as a metric for rhythm similarity. To do this, locomotor activity and body temperature data were binned every hour for each day post infection. For every mouse, a linear regression was performed using the binned data for each day post infection against that mouse's average rhythm before infection (24 data points; the hourly average over 48-72 hours of monitoring). This resulted in a daily $R^2$ value which reflects how much host rhythms deviate during infections from their patterns when mice were healthy. $R^2$ varies between 0 and 1; an $R^2$ of 1 indicates that the rhythm on that particular day post infection is identical to that observed before infection. |

Supplementary Table S1. Definitions of terms from Methods.

| | Locomotor activity | | | Body temperature | | |
|---|---|---|---|---|---|---|
| | $R^2$ | AIC mod 1 (inc G) | AIC mod 2 (exc G) | $R^2$ | AIC mod 1 (inc G) | AIC mod 2 (exc G) |
| "Asymptomatic" | 0.49 | **2372.08 (df=11)** | **2454.19 (df=5)** | 0.63 | **408.49 (df=8)** | **410.56 (df=4)** |
| "Moderate" | 0.18 | **1990.20 (df=11)** | **2020.86 (df=5)** | 0.65 | **500.21 (df=11)** | **631.73 (df=5)** |
| "Severe" | 0.57 | **2100.10 (df=11)** | **2223.13 (df=5)** | 0.94 | **251.10 (df=11)** | **284.25 (df=5)** |
| "Recovery" | 0.43 | 2087.71 (df=7) | 2091.94 (df=5) | 0.73 | **216.28 (df=11)** | **215.80 (df=5)** |

Supplementary Table S2. Coefficient of determination ($R^2$) and comparing model fits including (AIC mod 1) or excluding (AIC mod 2) the genotype term. Genotype differences using sine and cosine model terms as indicated by lower AIC values in all segments of infection in bold, with the exception of locomotor activity during the "recovery" segment. Associated with Figure 3.

| | Segment | Genotype comparison | Locomotor activity | | | | Body temperature | | | |
|---|---|---|---|---|---|---|---|---|---|---|
| | | | Estimate | Std. Error | t value | P value | Estimate | Std. Error | t value | P value |
| Night | "Asymptomatic" | AJ - AS | **-4.43** | **1.26** | **-3.50** | **0.046** | -0.12 | 0.35 | -0.34 | >0.99 |
| | | AJ - DK | **-11.65** | **1.26** | **-9.22** | **<0.01** | -0.28 | 0.35 | -0.81 | 0.99 |
| | | AS - DK | **-7.22** | **1.26** | **-5.72** | **<0.001** | -0.16 | 0.35 | -0.47 | >0.99 |
| | "Moderate" | AJ - AS | -0.95 | 1.26 | -0.75 | 0.99 | 0.54 | 0.35 | 1.57 | 0.91 |
| | | AJ - DK | -1.53 | 1.26 | -1.21 | 0.98 | -0.53 | 0.35 | -1.53 | 0.92 |
| | | AS - DK | -0.58 | 1.26 | -0.46 | >0.99 | -1.07 | 0.35 | -3.10 | 0.12 |
| | "Severe" | AJ - AS | **-7.14** | **1.33** | **-5.36** | **<0.01** | **-1.95** | **0.37** | **-5.32** | **<0.001** |
| | | AJ - DK | **-10.16** | **1.33** | **-7.62** | **<0.01** | **-2.08** | **0.37** | **-5.71** | **<0.001** |
| | | AS - DK | -3.02 | 1.26 | -2.39 | 0.43 | -0.14 | 0.35 | -0.41 | >0.99 |
| | "Recovery" | AJ - AS | 1.98 | 1.44 | 1.38 | 0.96 | -0.22 | 0.40 | -0.55 | 0.99 |
| | | AJ - DK | 0.37 | 1.44 | 0.26 | >0.99 | -0.25 | 0.40 | -0.62 | 0.99 |
| | | AS - DK | -1.61 | 1.26 | -1.28 | 0.98 | -0.03 | 0.35 | -0.08 | >0.99 |
| Day | "Asymptomatic" | AJ - AS | 2.63 | 0.79 | 3.34 | 0.07 | 0.19 | 0.33 | 0.56 | >0.99 |
| | | AJ - DK | 2.29 | 0.79 | 2.91 | 0.17 | 0.14 | 0.33 | 0.43 | >0.99 |
| | | AS - DK | -0.33 | 0.79 | -0.42 | >0.99 | -0.04 | 0.33 | -0.13 | >0.99 |
| | "Moderate" | AJ - AS | 2.05 | 0.79 | 2.61 | 0.30 | 0.52 | 0.33 | 1.59 | 0.90 |
| | | AJ - DK | 1.47 | 0.79 | 1.86 | 0.76 | 0.90 | 0.33 | 2.73 | 0.24 |
| | | AS - DK | -0.58 | 0.79 | -0.74 | 0.99 | 0.38 | 0.33 | 1.15 | 0.99 |
| | "Severe" | AJ - AS | -0.63 | 0.79 | -0.81 | 0.99 | **-1.76** | **0.33** | **-5.33** | **<0.01** |
| | | AJ - DK | -0.75 | 0.79 | -0.96 | 0.99 | **-2.00** | **0.33** | **-6.05** | **<0.01** |
| | | AS - DK | -0.12 | 0.79 | -0.15 | >0.99 | -0.24 | 0.33 | -0.72 | 0.99 |
| | "Recovery" | AJ - AS | 0.86 | 0.88 | 0.98 | 0.99 | 0.06 | 0.37 | 0.15 | >0.99 |

| | | | | | | | | | | |
|---|---|---|---|---|---|---|---|---|---|---|
| | | AJ - DK | 1.59 | 0.88 | 1.80 | 0.79 | 0.06 | 0.37 | 0.159 | >0.99 |
| | | AS - DK | 0.72 | 0.79 | 0.92 | 0.99 | 0.004 | 0.33 | 0.012 | >0.99 |

Supplementary Table S3. Genotypes differ in summary variables calculated for night and day during “asymptomatic” and “severe” segments. Each genotype is compared within each infection segment during night (ZT14-22) and day (ZT2-10). Reported are the parameter estimates, standard errors, t values for each comparison and adjusted p values (corrected for multiple comparisons). Significant differences between genotypes are highlighted in bold. Associated with Figure 3.

| Genotype comparison | Locomotor activity | | | | Body temperature | | | |
|---|---|---|---|---|---|---|---|---|
| | Estimate | Std. Error | t value | P value | Estimate | Std. Error | t value | P value |
| AJ - AS | **-0.18** | **0.02** | **-7.85** | **<1e-04** | **-0.20** | **0.04** | **-5.71** | **<1e-05** |
| AJ - DK | **-0.22** | **0.02** | **-9.42** | **<1e-04** | **-0.19** | **0.04** | **-5.31** | **<1e-05** |
| AS - DK | -0.04 | 0.02 | -1.68 | 0.22 | 0.01 | 0.03 | 0.42 | 0.91 |
| Genotype comparison | Red blood cell density | | | | Parasite density | | | |
| | Estimate | Std. Error | t value | P value | Estimate | Std. Error | t value | P value |
| AJ - AS | -0.29 | 0.14 | -2.01 | 0.11 | **24.94** | **5.41** | **4.61** | **<1e-04** |
| AJ - DK | **-0.75** | **0.14** | **-5.24** | **<1e-04** | **26.37** | **5.52** | **4.77** | **<1e-04** |
| AS - DK | **-0.46** | **0.12** | **-3.98** | **0.0003** | 1.43 | 5.24 | 0.27 | 0.96 |

Supplementary Table S4. Genotypes differ across infections in locomotor activity and body temperature disruption, red blood cell density and parasite density. Locomotor activity and body temperature disruption is calculated by comparing locomotor activity and body temperature rhythms for each day post infection back to rhythms from before infection (see Supplementary Table S1: Change in locomotor activity and body temperature, $R^2$). Red blood cell and parasite density is measured every day post infection from mouse blood. Each genotype is compared and parameter estimates, standard errors, t values for each comparison and adjusted p values (corrected for multiple comparisons) are reported. Significant differences between genotypes are highlighted in bold. Associated with Figure 4.

| | Genotype comparison | Locomotor activity | | | | Body temperature | | | |
|---|---|---|---|---|---|---|---|---|---|
| | | Estimate | Std. Error | t value | P value | Estimate | Std. Error | t value | P value |
| RBC density | AJ - AS | **-0.16** | **0.03** | **-6.21** | **<1e-05** | **-0.15** | **0.04** | **-3.91** | **0.0004** |
| | AJ - DK | **-0.02** | **0.03** | **-6.59** | **<1e-05** | -0.09 | 0.04 | -2.33 | 0.05 |
| | AS - DK | -0.01 | 0.02 | -0.34 | 0.94 | 0.06 | 0.04 | 1.62 | 0.24 |
| Parasite density | AJ - AS | **-0.09** | **0.03** | **-2.64** | **0.02** | -0.08 | 0.05 | -1.48 | 0.30 |
| | AJ - DK | **-0.20** | **0.03** | **-6.01** | **<1e-04** | **-0.14** | **0.05** | **-2.77** | **0.018** |
| | AS - DK | **-0.11** | **0.03** | **-3.17** | **0.005** | -0.06 | 0.05 | -1.19 | 0.46 |

Supplementary Table S5. The relationship between red blood cell and parasite density and locomotor activity and body temperature differs between genotypes. Plotting locomotor activity or body temperature disruption (see Supplementary Table S1: Change in locomotor activity and body temperature, $R^2$) against red blood cell (RBC) or parasite density. Each genotype is compared and parameter estimates, standard errors, t values for each comparison and adjusted p values (corrected for multiple comparisons) are reported. Significant differences between genotypes are highlighted in bold. Associated with Figure 5.
